# Iterated language learning is shaped by a drive for optimizing lossy compression

**DOI:** 10.1101/2025.05.23.655237

**Authors:** Nathaniel Imel, Jennifer Culbertson, Simon Kirby, Noga Zaslavsky

## Abstract

It has recently been theorized that languages evolve under pressure to attain near-optimal lossy compression of meanings into words. While this theory has been supported by broad crosslinguistic empirical evidence, it remains largely unknown what cognitive mechanisms may drive the cultural evolution of language toward near-optimal semantic systems. Here, we address this open question by studying language evolution in the lab via iterated learning. Across two qualitatively different domains (colors and Shepard circles), we find that semantic systems evolve toward the theoretical limit of efficient lossy compression, and over time, converge to highly efficient systems. This provides direct evidence that adult learners may operate under a bias to maintain efficiently compressed semantic representations. Moreover, it demonstrates how this bias can be amplified by cultural transmission, leading to the evolution of information-theoretically optimal semantic systems.

## Introduction

There are infinitely many ways to carve up the world into semantic categories. Human languages vary widely within this enormous space, but at the same time, appear to arrive at highly constrained solutions. For example, while some languages, like English, distinguish between ‘blue,’ ‘green,’ and ‘brown,’ and other languages do not make these distinctions, all attested language encode in their vocabularies a distinction between ‘black’ and ‘white’ (Berlin & Kay, 1969).

One pressure that is believed to underlie this constrained cross-linguistic semantic variation is the need to communicate efficiently (Kemp, Xu, & Regier, 2018; Gibson et al., 2019). Most relevant to our work is the theoretical framework of Zaslavsky et al. (2018), which is grounded in the mathematical theory of lossy compression (Shannon, 1959; Berger, 1971) and argues that languages evolve under pressure to efficiently compress meanings into forms via a fundamental information-theoretic principle known as the Information Bottleneck (IB; Tishby et al., 1999). This theory has been gaining broad empirical evidence across hundreds of languages and multiple semantic domains (Zaslavsky et al., 2018, 2019, 2021; Mollica et al., 2021). Its evolutionary predictions have also been directly supported by diachronic data documenting how a single language has changed over time (Zaslavsky et al., 2022). However, as noted by Levinson (2012), a major question remains open: What are the mechanisms by which languages become near-optimally efficient?

A promising approach for addressing this question is based on the iterated learning (IL) paradigm (Kirby et al., 2008), which offers a way to simulate the cultural evolution of language. However, prior work has so far yielded inconclusive answers. Carstensen et al. (2014) and Fedzechkina et al. (2012) have shown that IL can lead to semantic category systems with greater communicative informativeness. In contrast, others (Carr et al., 2020; Smith & Culbertson, 2018, 2025) have argued that pressure for minimizing complexity, which primarily supports learnability, rather than communicative informativeness, is the main driver of language evolution via IL. However, this literature has not adopted a consistent notion of efficiency, but rather has relied on differing formalizations of cognitive effort or informativeness.

Meanwhile, other researchers have begun to explicitly focus on how efficient compression in the IB sense might emerge. Chaabouni et al. (2021) and Tucker et al. (2022) showed how IB-efficient systems can emerge in artificial agents trained in communication games via reinforcement learning, without IL (see also Carlsson et al., 2021; Kågebäck et al., 2020), and Imel et al. (2024) showed a similar result with population dynamics instead of reinforcement learning. While these computational studies suggest that IL may not be crucial for the emergence of near-optimal semantic systems, their adaptive dynamics are nearly impossible to validate on actual human data. Carlsson et al. (2024) proposed a potential resolution by demonstrating that combining IL with communication can lead to efficient color naming systems. However, their results are based only on model simulations, rather than on IL with humans, and focus on the converged IL systems rather than on the full IL trajectories.

In this work, we aim to directly address Levinson (2012)’s question by revisiting IL data from chains of human learners through the lens of the IB framework. Specifically, we focus on two qualitatively different and well-studied domains (Figure 1): (a) color, which is a naturalistic domain that has been studied via IL by Xu et al. (2013), allowing us to test the theory on natural categories with respect to the previously established IB color naming model of Zaslavsky et al. (2018); and (b) Shepard circles (Shepard, 1964), which is a synthetic domain that has been studied via IL by Carr et al. (2020), allowing us to further test the theory while avoiding potential biases toward semantic categories in the participants’ native language. Across both domains, we find that learned semantic systems move toward the IB theoretical limit of efficiency, and over time, converge to highly efficient systems. Interestingly, the IL data we consider was collected without communication between participants, raising the possibility that adult learners may have an intrinsic bias toward maintaining efficient compression in their lexicon– a suggestion we return to in the discussion (see also (Fedzechkina et al., 2012; Smith & Culbertson, 2025)). Regardless of its origins, our work demonstrates how this tendency can be amplified by cultural transmission, leading to the evolution of IB-optimal systems.

**Figure 1:**
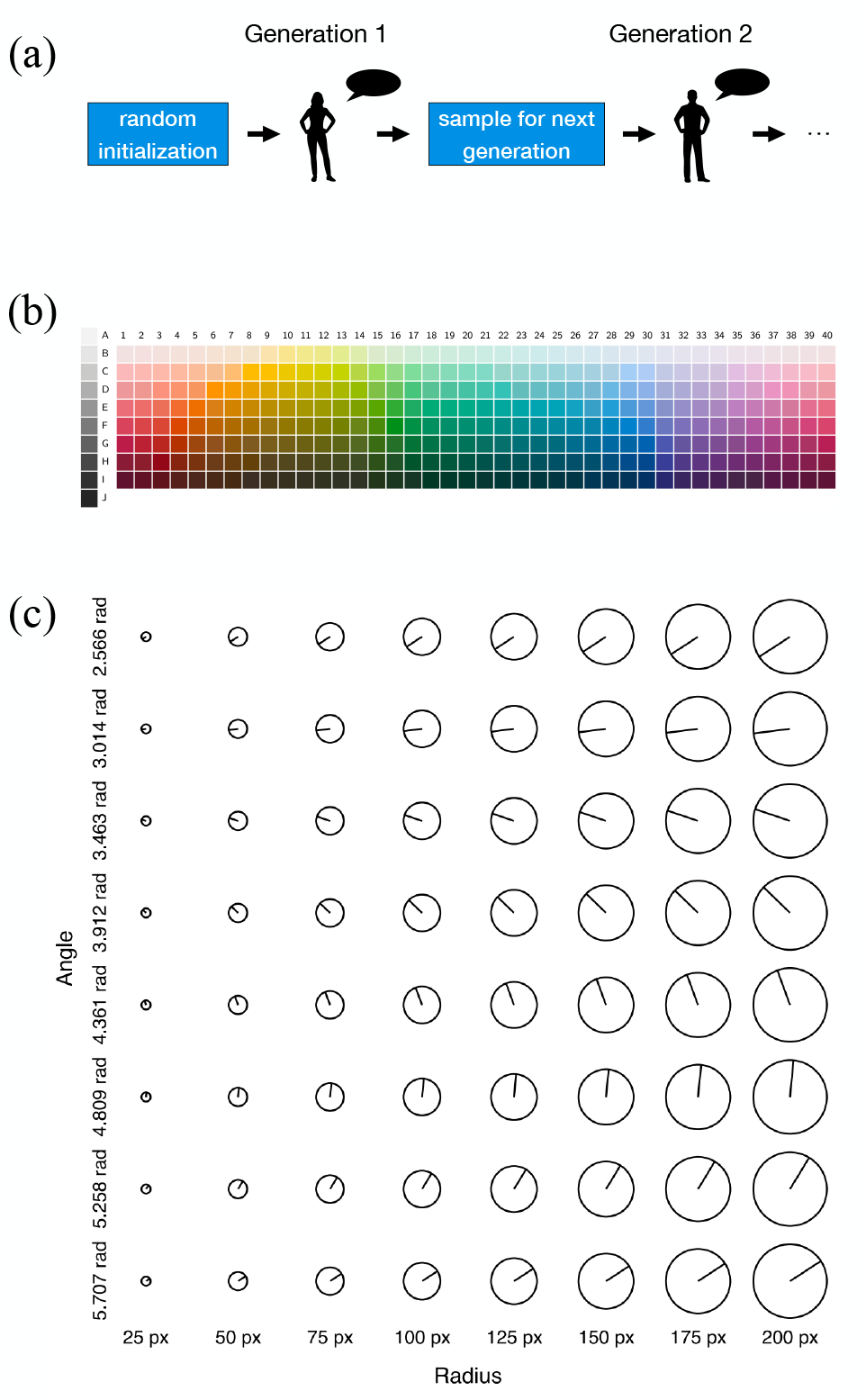
**(a)** Illustration of the iterated language learning experimental paradigm. **(b)** The WCS color naming grid, used as the stimuli in Xu et al. (2013)’s IL experiment. **(c)** Shepard circles stimuli grid, used in Carr et al. (2020)’s IL experiment.

### Language evolution via iterated learning

In a typical iterated learning (IL) experiment (Kirby et al., 2008), one participant is exposed to a set of data or stimuli and produces responses, which then serve as input for the next participant, forming a transmission chain (Figure 1a). In the experiments we examine here, participants are shown stimuli from a meaning space and asked to name them using one of a small set of made up words. This process is repeated until a complete category system is obtained from each participant. The next generation (a new participant) observes a limited sample of stimulus-word pairs generated from the previous generation’s system, and after training on these incomplete observations, the participant must reconstruct a full category system. This forced generalization from limited data introduces room for inference driven by inductive biases. As the literature on IL has shown, such generalization biases can strongly shape the resulting stationary distribution of languages that emerge (Griffiths & Kalish, 2007; Kirby et al., 2008). Intuitively, if language learning is a strong pressure shaping actual language evolution, then cognitive biases that operate during learning would restrict the emergent typological variation. This property of the IL paradigm makes it particularly useful for testing whether language evolution, as characterized by IL in the lab, is shaped by an intrinsic human bias toward efficient compression.

### Efficient compression and semantic systems

Our work builds on Zaslavsky et al. (2018)’s theoretical framework for the evolution of semantic systems, which is grounded in the mathematical theory of lossy compression (Shannon, 1959; Berger, 1971), and more specifically, in the Information Bottleneck (IB) principle (Tishby et al., 1999). This framework predicts that languages evolve to efficiently compress meanings into forms by optimizing the IB principle, which can be interpreted as striving to achieve an optimal tradeoff between the informational complexity and communicative accuracy of the lexicon. As the central components of this framework are not specific to any particular semantic domain, we first explain them in a domain-general way, as well as our application of this framework to iterated language learning. In the next sections we present our specific instantiations of this framework in two domains.

### Communication model

The framework is based on a basic communication model (Figure 2) that considers an inventory of signals 𝒲 (e.g., linguistic forms or artificial symbols) to communicate about a space of meanings, ℳ. Meanings are assumed to be mental representations or beliefs over world states 𝒰, formally defined as a probability distribution *m*(*u*) over world states *u* ∈ 𝒰. Meanings are drawn from an information source, *p*(*m*), which characterizes how often each meaning needs to be communicated. Given a meaning *m* ∼ *p*(*m*), a speaker produces a signal *w* using a stochastic production policy, also called an encoder, *q*(*w* | *m*), and then a listener interprets the signal by reconstructing an estimated belief state 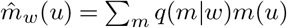. Note that this form of listener interpretations corresponds to a Bayesian listener. While this form is assumed here for simplicity, it is not an actual assumption but rather a derivation from the theory (see the SI of Zaslavsky et al., 2018).

**Figure 2:**
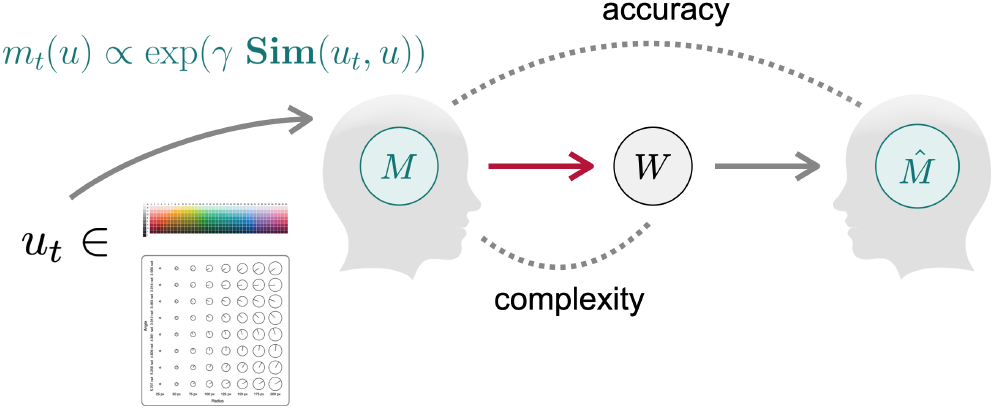
The communication model of Zaslavsky et al. (2018), in which meanings are grounded in an underlying representation of the environment captured by a similarity space (see main text). Here, we consider instantiations of this model in two domains: color and Shepard circles.

### Theoretical bound

A semantic system in this framework corresponds to an encoder, *q*(*w* | *m*), which maps meanings to signals. An optimal semantic system, according to the IB principle, is an encoder that satisfies a tradeoff between its informational complexity and communicative accuracy. Complexity, also known as information rate (Cover & Thomas, 2006), is defined by the mutual information between speaker meanings and signals,

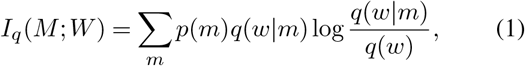

which captures the number of bits, on average, that is required to encode meanings with signals. Maintaining low complexity corresponds to using fewer bits for communication, i.e., achieving high compression rate, which in turn, can be translated to affording a smaller lexicon size. For example, minimal complexity can be achieved by compressing all possible meanings into a single signal. This, however, will result in very poor accuracy. Accuracy, in IB terms, is quantified by

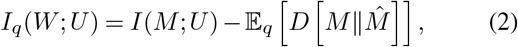

which corresponds to maintaining a language that is informative about the speaker’s intentions. Maximizing accuracy, as expressed in Equation (2), amounts to minimizing the expected distortion between speaker and listener meanings, i.e., the Kullback-Leibler (KL) divergence 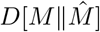. We say that a language’s semantic system is *efficient* to the extent that its encoder *q* minimizes the IB tradeoff:

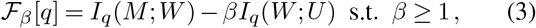

where *β* is a free parameter controlling the tradeoff between pressure to minimize complexity and pressure to maximize accuracy. The solutions to this optimization problem define the IB theoretical limit of efficiency, which means that no system can lie above this theoretical bound (see Figure 4).

### Quantitative predictions for iterated learning

Zaslavsky et al. (2018) theorized that languages evolve under pressure to optimize the IB tradeoff and derived from this theory two quantitative predictions: (i) semantic systems of actual languages should lie near the IB theoretical bound, namely, their deviation from optimality should be small; (ii) the optimal IB encoders along the bound should resemble actual semantic systems observed across languages, i.e., there should be high alignment between human and IB encoders. The latter can be quantified by the Normalized Information Distance (NID; Kraskov et al., 2005; Vinh et al., 2010) between languages and their nearest optimal IB encoders. These predictions have been supported empirically across hundreds of languages and multiple domains, including color (Zaslavsky et al., 2018, 2022), pronouns (Zaslavsky et al., 2021), containers and animal taxonomies (Zaslavsky et al., 2019), and grammatical number, tense, and evidentiality (Mollica et al., 2021).

While this broad empirical evidence suggests that semantic systems across languages achieve near-optimal compression via IB, it remains largely unknown what mechanism could drive populations toward optimality. One possibility is that a drive for maintaining efficiently compressed representations operates in individuals. If this is true, then such a bias should be amplified by chains of iterated language learning, leading these chains toward nontrivially efficient systems that lie near the IB bound. Formally, this means that (i) systems should converge to intermediate complexity-accuracy trade-offs and (ii) we would expect that across generations of iterated learning, the deviation from optimality will tend to decrease and the human-IB alignment will tend to increase, eventually converging to highly efficient systems.

## Studies

To test these theoretical predictions, we turn to two empirical studies, one in the domain of color and one in a synthetically-generated geometric domain. In each study, we evaluate IL human data with respect to an IB model instantiation. In this section we describe the data and model instantiations, and in the next section we discuss our results.

### Study 1: Color naming

Color naming is an important testbed for analyzing efficiency in IL for three main reasons. First, it is well-documented across languages, with the World Color Survey (WCS; Cook, Kay, & Regier, 2005) serving as one of the most comprehensive and influential collections of cross-linguistic semantic category data (Berlin & Kay, 1969; Kay et al., 2009). Second, this domain has previously been studied via IL by Xu et al. (2013), providing human IL color naming data with respect to the WCS color grid. Third, Zaslavsky et al. (2018) applied the IB principle in this domain, yielding an IB color naming model that has gained extensive empirical support on the WCS data as well as on more recent diachronic data documenting language change (Zaslavsky et al., 2022). Taken together, color naming is a rare case where we have a combination of a well-established IB model and human IL data.

#### IL data

The empirical basis for our color naming study is the IL data from Xu et al. (2013). In their experiment, participants were asked to learn and transmit novel systems of color terms across 13 generations. We focus on their main results that include twenty iterated learning chains (a sample shown in Figure 3a), each initialized with a random partition of the WCS grid. These chains vary in the number of allowed color terms, ranging from two to six. Participants were shown a set of randomly selected colors (with each term having an equal number of instances), generated uniquely for each chain, and paired with corresponding pseudo words. After training, participants were asked to label all 330 colors of the WCS grid. Xu et al. (2013) found that over time, the IL chains become increasingly regular and resembling the color naming systems documented in the WCS dataset. This suggests that iterated language learning in the lab evolves captures some cognitive biases and constraints that may lead to attested structures in color naming across languages.

**Figure 3:**
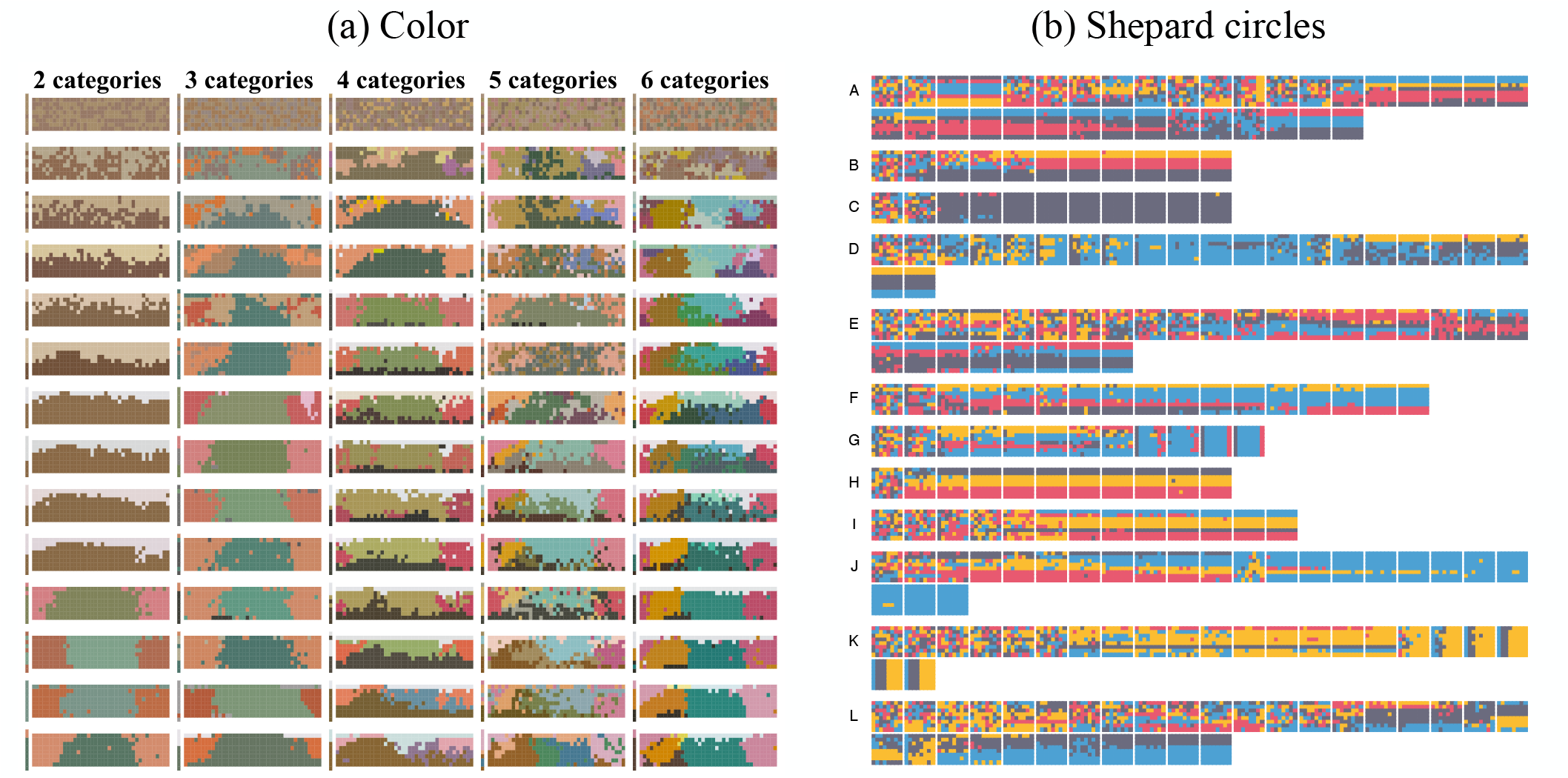
**(a)** Sample of IL chains for color naming (data from Xu et al., 2013, figure from Griffiths & Zaslavsky, 2020). Columns correspond to chains with *K* = 2, 3, 4, 5, 6 categories; rows correspond to generations. Each system is shown as mode map over the WCS grid, where each color chip is colored by its label and each label is associated with the average color of its category. **(b)** All IL chains for naming Shepard circles (data and figure from Carr et al., 2020). Rows (A-L) correspond to individual chains. Each system is plotted over the stimuli grid of Figure 1c, where colors correspond to unique labels.

#### IB model instantiation

We base our efficiency analysis on the previously published IB color naming model of Zaslavsky et al. (2018).^1^ In this model, the set of world states 𝒰 is taken to be the 330 color chips from the WCS grid, and meanings over colors are grounded in the CIELAB perceptual space such that each target color referent, *u*_*t*_ ∈ 𝒰, is represented as a Gaussian distribution *m*_*t*_(*u*) centered around *u*_*t*_. The need distribution over meanings in this model, *p*(*m*), was estimated using the method of least-informative priors (see Zaslavsky et al., 2020 for an extensive evaluation of this need distribution).

### Study 2: Shepard circles

While color provides a crucial test case in our context, one important limitation of this domain is that color categories are also encoded in the participant’s languages, and therefore the data might reflect a bias toward the color categories that participants already know and use, rather than evolve via cultural transmission. To address this potential concern, we consider a second domain, based on Shepard circles (Shepard, 1964), which was studied in this context by Carr et al. (2020). In contrast to color, this domain is based on an abstract synthetically-generated geometric stimulus set spanned by two features, radius and angle, with eight discrete values for each feature, yielding the stimulus grid shown in Figure 1c.

#### IL data

We base our efficiency analysis of this domain on data from the iterated language learning experiment of Carr et al. (2020).^2^ In their study, they simulated 12 IL chains (labeled A–L in Figure 3b). Participants were restricted to using exactly four labels, and each chain was randomly initialized using these four labels. In the training phase, participants were shown a random, representative sample of half of the 64 stimuli, and in the production phase they were asked to label the entire stimulus set. Chains were simulated until a convergence criterion applied when two consecutive participants produced identical languages, yielding 11–35 generations.

#### IB model instantiation

In order to apply the IB framework to the domain of Shepard circles, we first need to instantiate the communication model (Figure 2) for this domain. This requires specifying two components: (i) an underlying representation of the domain in which meanings are grounded, and (ii) a need distribution over meanings. For the latter, we simply take the uniform distribution. To specify the space of grounded meaning representations, we first define the set of world states 𝒰 ⊂ ℝ ^2^, such that each element in the domain *u* ∈ 𝒰 can be represented as *u* = (*a*_*u*_, *r*_*u*_) where *r*_*u*_ and *a*_*u*_ are the values of its radius and angle respectively. Following prior work (e.g., Xu et al., 2016; Zaslavsky et al., 2022), we assume that each meaning takes the form of a similarity-based distribution, i.e., *m*_*t*_(*u*) ∝ exp(− *γ* · *d*(*u, u*_*t*_)). While it may seem natural at first to take *d*(*u, u*_*t*_) to be the Euclidean distance, Shepard (1964) argued in his foundational work that human perceptual similarities in this domain are inconsistent with a Euclidean distance. Instead, people tend to attend to only a single feature dimension when categorizing stimuli. This effect is strikingly visible in the IL chain data (Figure 3b): participants do not converge to systems that partition the space along both dimensions (e.g., into four quadrants). Instead, eight chains converge to systems that distinguish only along angle and two to systems distinguishing only radius (the remaining two converge to degenerate systems which do not make any semantic distinctions).

To capture Shepard’s observation, we consider a Mahalanobis distance, a weighted generalization of Euclidean distance. To this end, we fix *γ* using the domain-agnostic method from (Eisape et al., 2020; Zaslavsky et al., 2021), and incorporate a feature-attention parameter *α* ∈ [0, 1]:

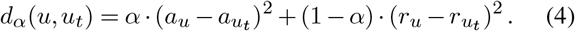

When *α* = 1 the model attends only to angle and when *α* = 0 it attends only to radius. While this parameter can vary across participants and over time, we assume for simplicity that it is fixed. Because we do not know what is the right value of *α*, we ‘reverse-engineer’ it by gridsearching 11 values, recording for each chain the value that led to lowest deviation from optimality. Notably, *α* = 0.7 emerged as the best-fitting value for six of the twelve chains, and *α* = 0.6 for one. This aligns with the observed angle bias in the data. In chains that shift attention to radius, the best-fit values were *α* = 0.3 or 0.4, demonstrating that our approach can effectively identify *α*.

Based on these findings, we fix *α* = 0.7. While it may seem that this approach is favorable to our model, note that because *α* is fixed across all generations and chains, it cannot account for any efficiency trends either toward or away from the theoretical bound. In addition, it does not imply that any of the IL systems, even those that were fitted with this value, are actually significantly efficient. Having said that, we wish to highlight that this model instantiation is only an initial exploration, and in future work we intend to ground our model in non-linguistic similarity judgments, as has been done in other domains (e.g., Xu et al., 2016; Zaslavsky et al., 2019).

## Results

Given the IL data and IB model instantiations discussed above, we are ready to test whether IL chains are shaped by pressure to efficiently compress meanings into words. To this end, for each system along each chain, we evaluated its complexity (Eq. 1), accuracy (Eq. 2), and deviation from optimality defined by 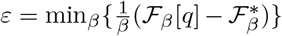 where 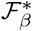 is the optimal value of ℱ_*β*_ (see Zaslavsky et al., 2018, for more details). We also computed a measure of system’s ‘Human-IB’ alignment’, operationalized as the Normalized Information Distance (Vinh et al., 2010) between the system and its *ϵ*− fitted IB system. Figure 4 shows the human IL complexity-accuracy trajectories (colored curves) on the IB information plane. In both domains, the trajectories tend to approach the IB theoretical bound of efficiency (black curve). Unsurprisingly, these trajectories are rather noisy, and some do not reach the bound, such as chains K and G in Figure 4b, which converged to a radius-based rather than angle-based solution. However, there is a clear emergent tendency to approach the theoretical bound. To validate this observation quantitatively, Figure 5a-b shows that across all chains in both domains, the IL systems become more efficient over time (deviation from optimality decreases) and more similar to the IB-optimal encoders (human-IB alignment increases).

**Figure 4:**
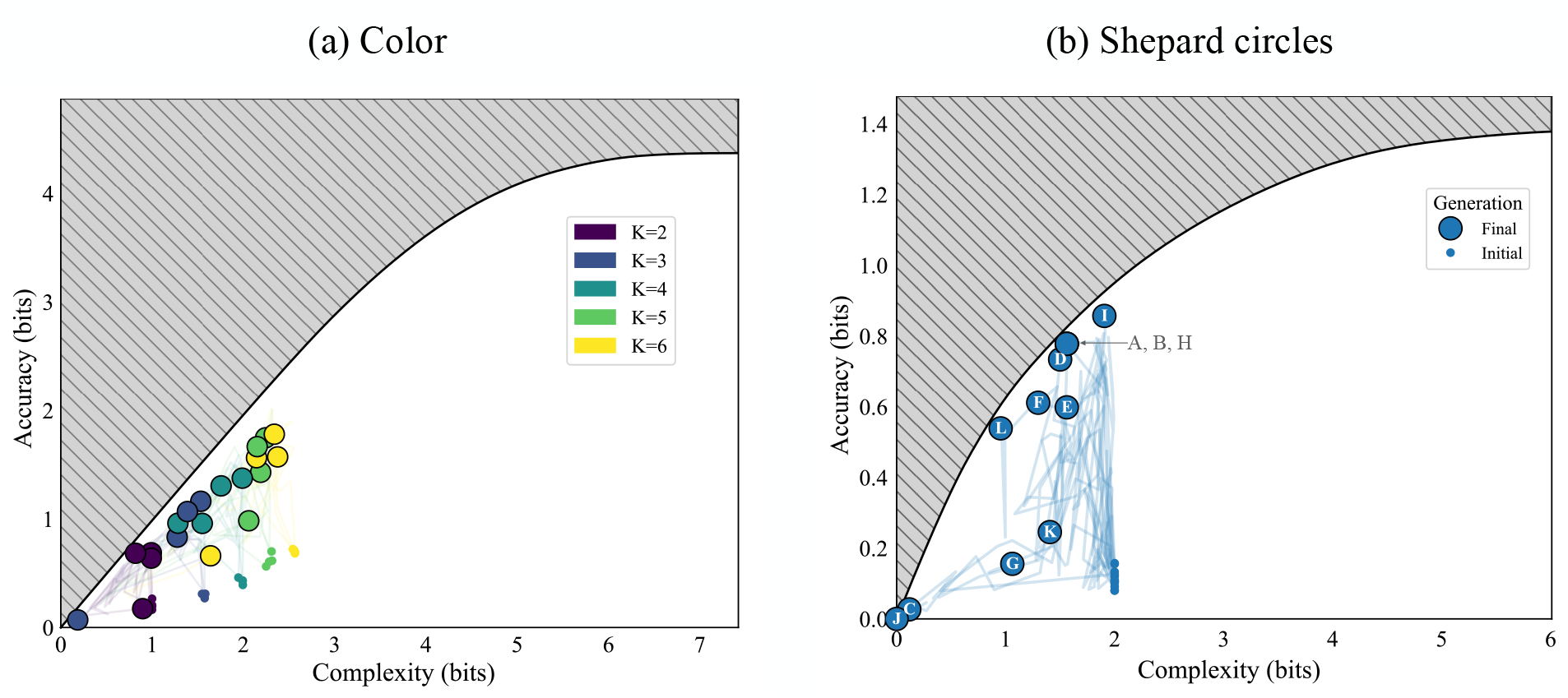
Iterated learning trajectories on the IB plane. Initial random naming systems (small colored dots) evolve in non-monotonic trajectories (colored curves) towards the IB bound (black curve). Larger colored dots mark final generations and are labeled by chain, as in Figure 3.

**Figure 5:**
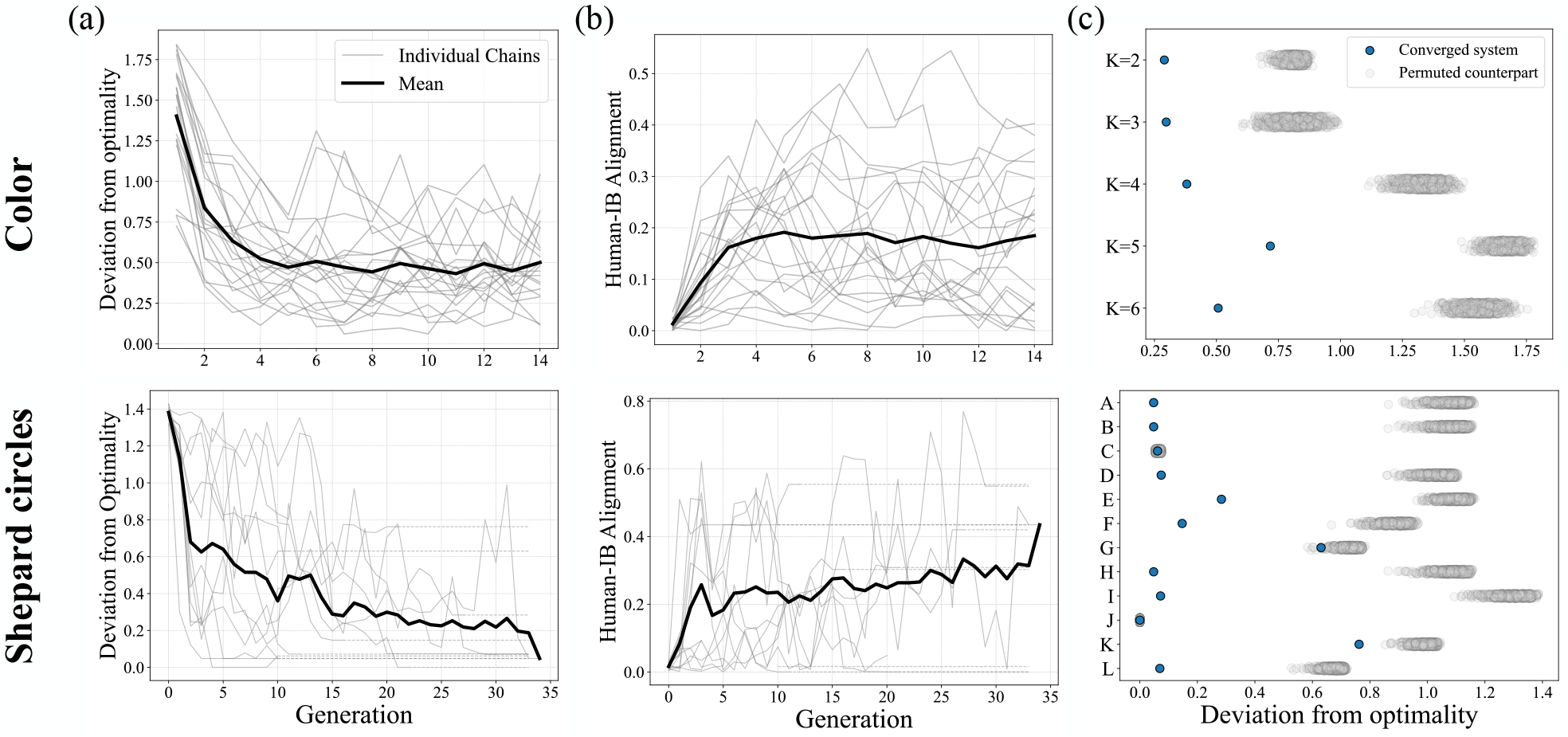
Quantitative analysis of IL chains in the domain of color (top) and Shepard circles (bottom). (a) Deviation from optimality across generations. Thin dashed lines reflect converged chains with padding to match the maximal length. (b) Similarity of IL systems to IB optima across generations. (c) Non-degenerate IL systems are more efficient than their permuted counterparts. For color, only the chains from Figure 3a are visualized, and similar results hold across chains.

Although the IL chains approach the bound, they do not reach it fully. To test whether the IL chains nevertheless converge to significantly near-optimal systems, we compared the efficiency of the final generation of each chain with 1, 000 hypothetical counterparts generated by permuting the generation’s system (Figure 5c). In all but two chains from each study, converged systems were more efficient than all of their permuted counterparts. Crucially, this suggests that participants do not operate under one pressure alone, such as only minimizing complexity, but rather gravitate toward efficiently balancing the IB complexity-accuracy tradeoff.

## Discussion

We set out to identify a cultural evolutionary mechanism that could explain how human semantic systems converge to near-optimal IB solutions. We revisited two prominent IL studies through the lens of IB and found that adult learners restructure category systems toward highly efficient solutions. This suggests that cultural transmission via IL may play a key role in driving human languages toward near-optimally efficient semantic systems. Notably, most chains did not converge to the most informative (and most complex) solution for their lexicon size nor to the simplest possible (non-informative) solution. Instead, they converged to non-trivial trade-offs.

This finding is somewhat surprising given that there was no communication between participants. That is, learnability, rather than communication, is the main cultural mechanism driving these chains. Intuitively, one might expect that learnability alone would drive the chains toward non-informative solutions, which are easiest to learn (Kirby et al., 2015; Smith & Culbertson, 2018, 2025; Carr et al., 2020; Carlsson et al., 2024). One possible explanation for why we do not observe this in the data is that perhaps people have a flexible intrinsic bias toward informativity. This bias might be weaker without an explicit need to communicate, and more generally, can be beneficial for social agents (Tucker et al., 2022, 2025). An alternative explanation is that perhaps this observation simply reflects an artifact of the experimental design. Therefore, an important direction for future research is to tease apart the influence of learnability and communication in our context. Either way, we predict that adding an explicit pressure for communication will drive the chains further toward the IB theoretical bound, and possibly also higher along the bound.

## Acknowledgments

NZ is grateful to Jing Xu for sharing the iterated color naming data and for helpful discussions about it.

## Footnotes

1 Using publicly available model and accompanying code from https://github.com/nogazs/ib-color-naming

2 Using publicly available data and accompanying code from https://github.com/jwcarr/shepard.

